# Efficacy of Tumor Treating Fields (TTFields) in triple negative breast cancer cells and synergy with PARP inhibitor talazoparib

**DOI:** 10.64898/2026.10.09.757989

**Authors:** Maryam Ghandali, Praveen Srinivasan, Shengliang Zhang, Yutong Xia, Eric T. Wong, Clark C. Chen, Robert W. Sobol, Stephanie L. Graff, Lanlan Zhou, Benedito A. Carneiro, Yaara Porat, Moshe Giladi, Naama Flint-Brodsly, Wafik S. El-Deiry

## Abstract

Breast cancer is the most diagnosed cancer worldwide. Triple-negative breast cancer (TNBC) is a subtype with a poor prognosis because of its aggressiveness and high likelihood of relapse, making its treatment particularly challenging. We explored the effect of Tumor Treating Fields (TTFields) as a novel treatment option for breast cancer, which can be used together with other therapies, such as PARP inhibitors. TTFields use alternating electric fields to inhibit cell growth and induce cellular damage. Our findings indicate that TTFields increase expression of the universal cell cycle CDK inhibitory protein p21(WAF1)/CDKN1A independently of *TP53* mutational status. Moreover, when used together with the PARP inhibitor talazoparib, the combination further upregulates p21 protein. Both TTFields alone and together with talazoparib induce cell death as documented by PARP cleavage while reducing markers of cell growth and anti-apoptotic proteins, including p-STAT3, STAT3, and MCL-1. Additionally, TTFields together with talazoparib induce DNA damage, as indicated by increased γH2AX expression. The combination of talazoparib plus TTFields treatment decrease PDL-1 levels and given its utility in predicting response to PD-1/PD-L1 by mediating immune evasion, this raises the possibility of future combination for immune modulation in cancer therapy. Our preclinical results suggest TTFields alone or together with PARP inhibitors such as talazoparib as a promising therapeutic approach for further development for the treatment of breast cancer.

## Introduction

Breast cancer is the most diagnosed cancer worldwide (1). In the United States, an estimated 313,510 new cases of breast cancer and 42,780 breast cancer-related deaths were expected in 2024. Among women, an estimated 310,720 new cases and 42,250 deaths were expected. (2). Triple negative breast cancer is an aggressive subtype known by higher likelihood of metastasis, recurrence and poor prognosis (3). The standard treatment of triple negative breast cancer (TNBC) includes surgery, radiotherapy and adjuvant chemotherapy (4). Because of the aggressiveness and high relapse rate of TNBC, new therapeutic alternatives have been developed (5).

About 25% of patients with triple-negative breast cancer harbor *BRCA1* or *BRCA2* mutations, which can make them more sensitive to PARP inhibitors (6). BRCA1 and BRCA2 are involved in homologous recombination, an important pathway for repairing DNA double-strand breaks. When BRCA1 or BRCA2 is mutated, this repair pathway is impaired, making cancer cells more dependent on other DNA repair mechanisms for survival. PARP inhibitors block one of these alternative repair mechanisms, leading to accumulation of DNA damage and eventually cell death. This is known as synthetic lethality, where loss of either repair pathway alone may not cause cell death, but impairment of both pathways becomes lethal to the cancer cell (7). talazoparib is a PARP inhibitor approved for the treatment of HER2-negative, locally advanced or metastatic breast cancer with germline *BRCA1/2* mutations (8).

Tumor treating fields (TTFields) use electric fields to disrupt cell division and cause cell death. The electric fields have a frequency between 100-500 khz (9). TTFields are FDA-approved for treatment of Glioblastoma multiforme (GBM), malignant pleural mesothelioma (MPM), non-small lung cancer and pancreatic cancer (10). Previous studies showed that TTFields can induce “BRCAness” and therefore increased vulnerability to DNA-damaging agents (11,12,13,14,15,16). STAT (signal transducer and activator of transcription) proteins are key transcription factors that become activated in response to growth factors and cytokines. Activation of STATs in tumors can cause chemoresistance and poor patient survival (17).

As shown before, TTFields influence different molecular pathways, reducing markers of cell growth and anti-apoptotic proteins and increasing markers of DNA damage (18). The combination of TTFields with sorafenib in colorectal cancer showed downregulation of STAT3 (19). Another study demonstrated that the combination of TTFields with hyperthermia can cause downregulation of STAT3 (20).

In breast cancer, various secreted factors like IL-6, IL-10 activate STAT3, and these factors are linked to advanced cancer and poor prognosis (21). However, the role of PARP inhibitors on STAT3 remains controversial. While olaparib has been shown to increase STAT3 activation, Niraparib inhibits its activity (22, 23).

We investigated the effect of TTFields, talazoparib or their combination on STAT3 and other anti-apoptotic proteins associated with its transcriptional activity. To explore common pathways, we evaluated markers of cell death, cell cycle and DNA damage like p53, p21 and γH2AX following exposure of TNBC cells to TTFields, talazoparib or their combination. Our results offer new insights into the molecular pathways affected by TTFields in breast cancer and suggest a potential new combination therapy for its treatment.

## Material and Methods

The human TNBC cell lines MDA-MB-436 and MDA-MB-231 were cultured in DMEM medium plus 10% fetal bovine serum and 1% penicillin-streptomycin. The MDA-MB-468 cells were cultured in RPMI 1640 medium with 10% fetal bovine serum, and 1% penicillin-streptomycin, The cells were cultured at 37°C.

### *In vitro* application of Tumor Treating Fields (TTFields)

Cells were seeded directly into inovitro™ ceramic dishes at densities adjusted according to the cell line and treatment duration to achieve an appropriate confluence at the experimental endpoint. Following cell attachment, the dishes were placed on inovitro™ base plates connected to a TTFields generator (Novocure, Haifa, Israel). TTFields were applied at a frequency of 150 kHz, with a target temperature of 37°C maintained within the ceramic dishes throughout treatment. Control cells were cultured in identical inovitro™ ceramic dishes under standard incubator conditions without TTFields exposure. Cells were exposed to TTFields for the indicated durations and subsequently collected for downstream analyses.

### Immunoblotting

After the specified incubation period, the cells were lysed using a protein lysis buffer. The BCA protein assay kit (Thermo Fisher Scientific, Carlsbad, CA, USA) was used to measure protein concentration. Equal protein amounts were loaded onto a NuPAGE 4-12% Bis-Tris gel (Thermo Fisher Scientific), then transferred onto PVDF membranes. the membrane was blocked with 5% non-fat milk. The membranes were incubated overnight at 4°C with primary antibodies in 5% non-fat milk. The next day, the membranes were washed and treated with the appropriate secondary antibodies in 5% non-fat milk. The primary antibodies used were cleaved PARP, γH2AX, MCL-1, phospho-STAT3, STAT3, vinculin from Cell Signaling; anti-p21 antibody from Invitrogen; anti-p53 antibody from Santa Cruz Biotechnology; and Ran antibody from BD Biosciences. Anti-Rabbit and anti-mouse antibodies were used as secondary antibody.

| Antibody | Catalog number | Company | Dilution |
| --- | --- | --- | --- |
| Cleaved PARP (Asp214) (19F4) | 9546 | Cell signaling | 1:1000 |
| Ran | 610341 | BD Transduction Laboratories | 1:5000 |
| Vinculin E1E9V | 13901 | Cell signaling | 1:1000 |
| Phospho-Histone H2A.X (Ser139) | 2577 | Cell signaling | 1:1000 |
| Phospho-Stat3 (Tyr705) (D3A7) | 9145 | Cell signaling | 1:1000 |
| Stat3 (124H6) | 9139 | Cell signaling | 1:1000 |
| MCL-1 | 4572 | Cell signaling | 1:1000 |
| PD-L1 (E1L3N®) XP® | 13684 | Cell signaling | 1:1000 |
| p53 (DO-1): sc-126 | 12817 | Santa cruz biotechnology | 1:500 |
| p21 (WAF1, Cip1) | 14671581 | Invitrogen | 1:500 |

## Results

### TTFields increase p21(WAF1) protein levels in TNBC cell lines independent of *TP53* mutational status

To explore possible induction of DNA damage by TTFields and its effect on cell growth or cell cycle arrest, we investigated the p53-p21 pathway. Levels of the cyclin-dependent kinase (CDK) inhibitor p21(WAF1)/CDKN1A increase in response to multiple factors that induce cell-cycle arrest including DNA damage or other cellular stresses that activate p53. p21 can increase in both a p53-dependent and -independent manner (24). After TTFields single exposure, both MDA-MB-468 (*TP53*-mutant; homozygous *TP53* R273H, *BRCA1* wild-type) and MDA-MB-436 cells (*TP53* mutant; frameshift mutation p.Glu204fs*7 leading to truncated p53 protein and loss of function, *BRCA1* mutated; hemizygous splice donor mutation in exon 20 c.5277+1G>A leading to a C-terminal truncated BRCT domain protein) showed an increase in p21 protein expression (**Figure 1**; **Table 1**). This finding is consistent with previous studies demonstrating the effects of TTFields on p21 expression in MPM, ovarian and gastric cancer, as well as in pan-cancer models (11,12,13,14,15,16). We demonstrate that this upregulation of p21 is independent of *TP53* mutational status (**Figure 1**; **Table 1**).

**Figure 1.**
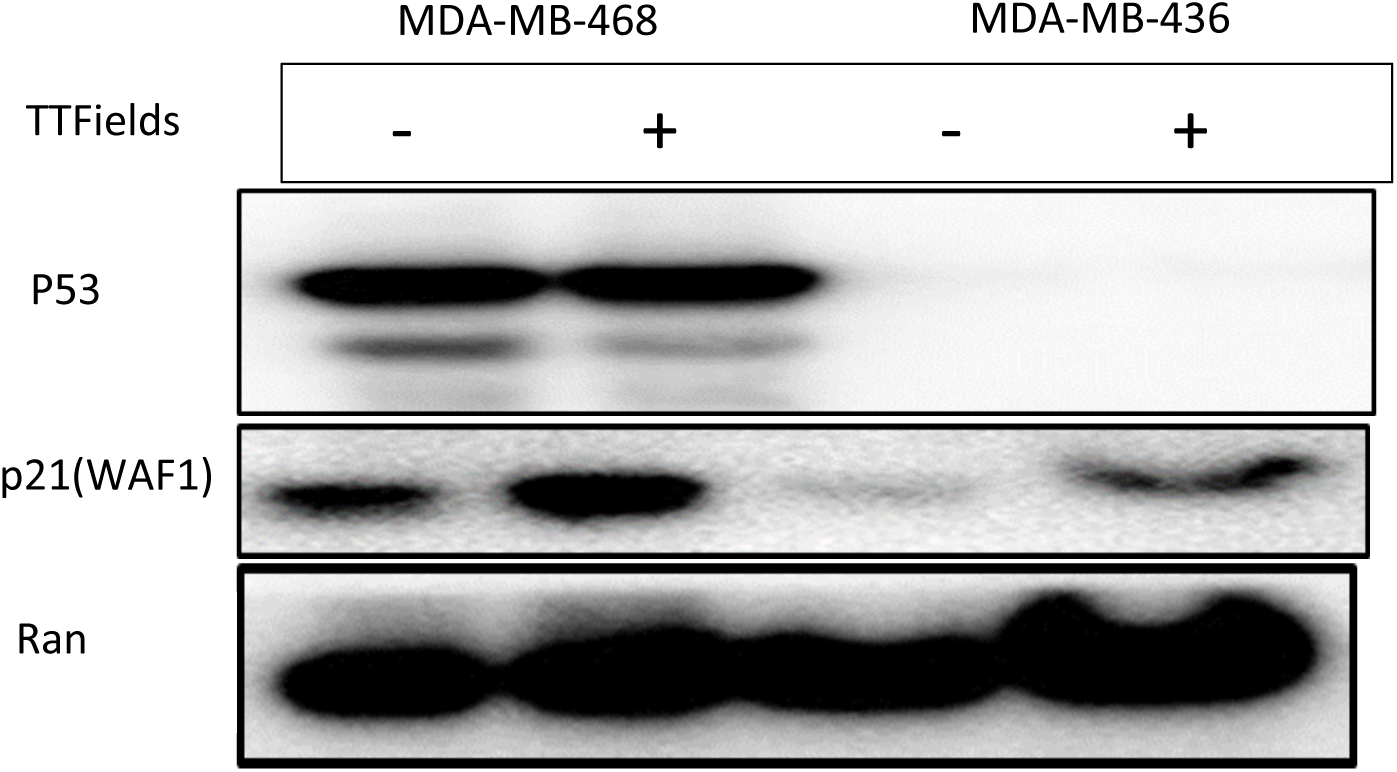
Upregulation of p21(WAF1)/CDKN1A protein by TTFields in human TNBC cells occurs independently of *TP53/BRCA1* mutational status. MDA-MB-468 (*TP53*-mutant; homozygous *TP53* R273H, *BRCA1* wild-type) and MDA-MB-436 cells (*TP53* mutant; frameshift mutation p.Glu204fs*7 leading to truncated p53 protein and loss of function, *BRCA1* mutated; hemizygous splice donor mutation in exon 20 c.5277+1G>A leading to a C-terminal truncated BRCT domain protein) were treated with TTFields (150 khz) for 24 hours. p21(WAF1)/CDKN1A) protein was upregulated independent of *TP53* mutational status.

**Table 1:**
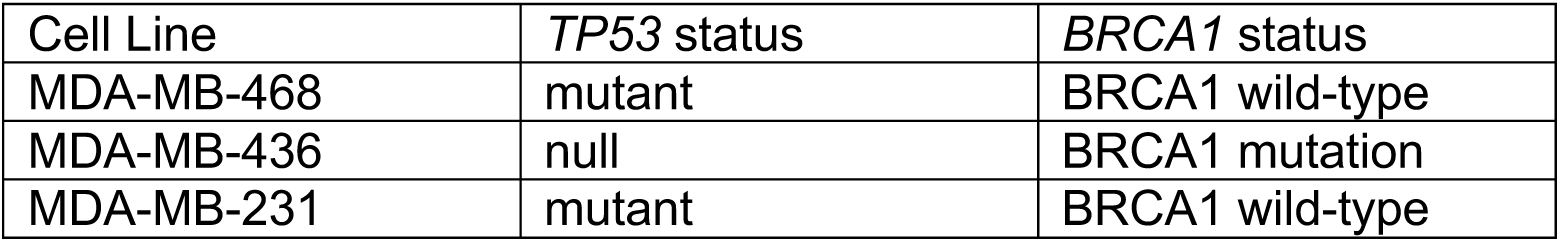
Human TNBC cell lines used in experiments and their known *TP53/BRCA1* mutational status.

| Cell Line | *TP53* status | *BRCA1* status |
| --- | --- | --- |
| MDA-MB-468 | mutant | BRCA1 wild-type |
| MDA-MB-436 | null | BRCA1 mutation |
| MDA-MB-231 | mutant | BRCA1 wild-type |

### TTFields together with PARP inhibitor talazoparib upregulate p21 protein in human TNBC cell lines

MDA-MB-468 (*TP53*-mutant; homozygous *TP53* R273H, *BRCA1* wild-type) and MDA-MB-231*(TP53* mutant; missense R280K *TP53* mutation in DNA-binding domain, *BRCA1* wild-type) cells were treated with TTFields and PARP inhibitor talazoparib for 24 hours. p21 protein levels were found to be increased as result of the combination of exposure to TTFields and PARP inhibitor talazoparib compared to control and each treatment alone in both cell lines (**Figure 2**). The PARP cleavage and γH2AX upregulation was heterogeneous among the TNBC cell lines. MDA-MB-231 cells showed more PARP cleavage (indicative of apoptosis) and γH2AX (indicative of DNA double strand breaks) following combination of TTFields and PARP inhibitor talazoparib as compared to single treatments. (**Figure 2B**). MDA-MB 468 cells showed PARP cleavage after treatment with both TTFields alone and combination of TTFields and Talazoparib. γH2AX appeared more upregulated because of TTFields treatment alone (**Figure 2A).** However, with longer treatment (72 hours) in MDA-MB-468 cell line, γH2AX upregulation following exposure to TTFields was decreased and TTFields together with talazoparib showed more DNA damage and cell death as demonstrated by PARP cleavage (**Figure 3B**).

**Figure 2.**
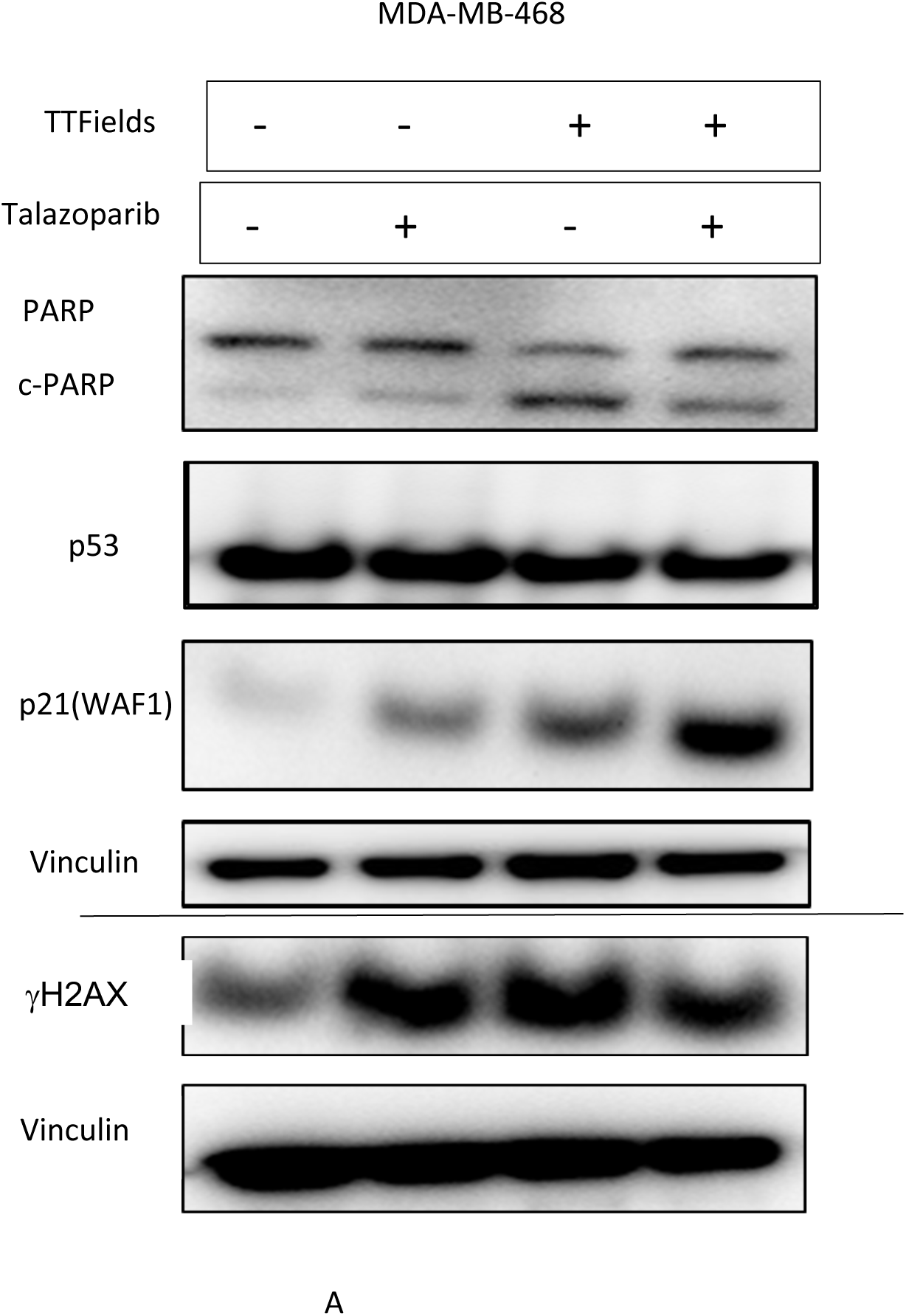
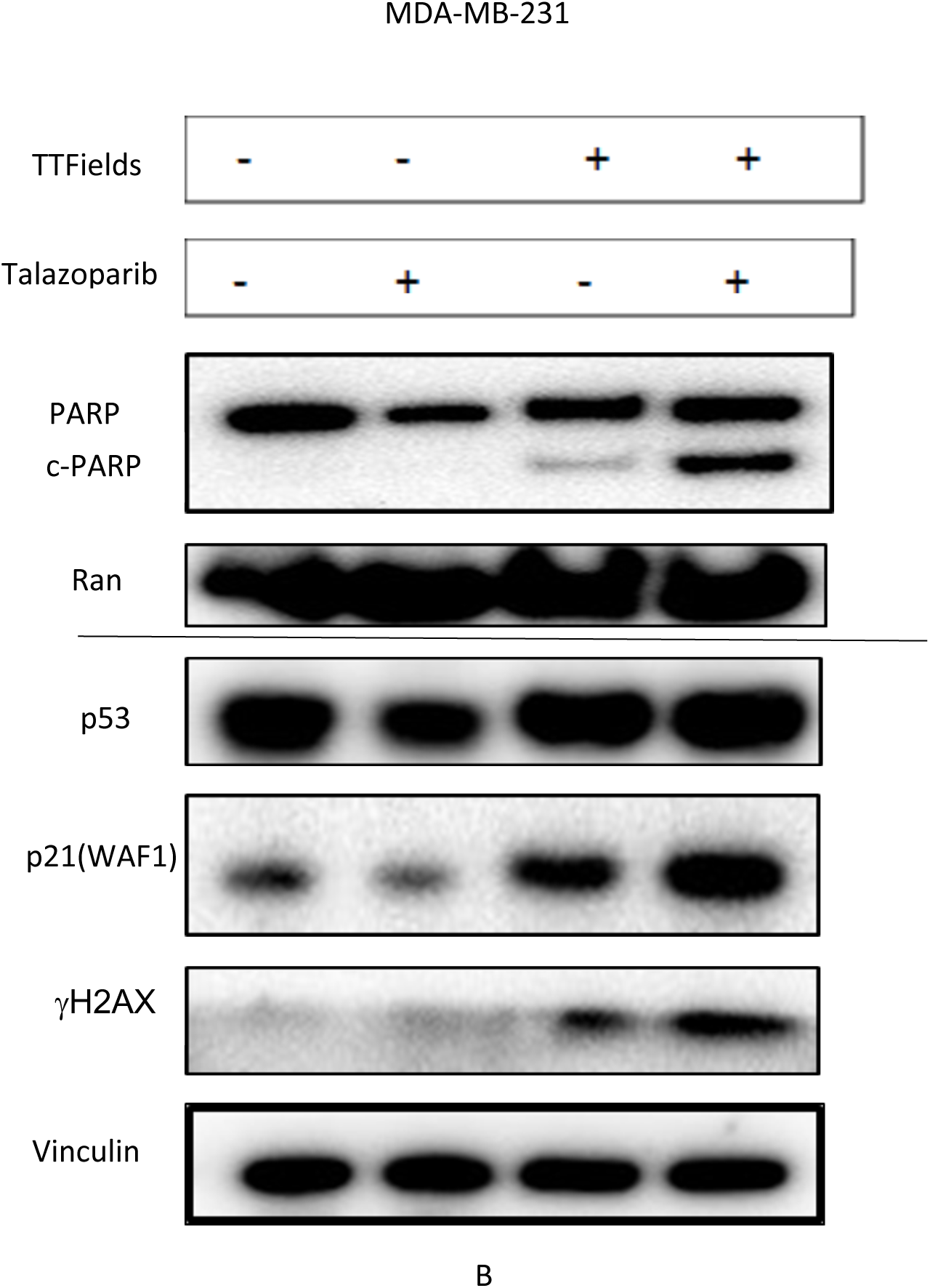
Induction of p21(WAF1), γH2AX, and cleaved PARP by TTFields -/+ talazaporib in human TNBC cells. **(A)** MDA-MB-468 cells (*BRCA1* wild-type, *TP53* mutated) were treated with either TTFields 150 khz, talazoparib at 2.5 µM or the combination for 24 hours, TTFields and combination with talazaporib caused PARP cleavage as compared to control. p21(WAF1) was upregulated in each treatment alone and by the combination. yH2AX was upregulated following each treatment. **(B)** MDA-MB-231*(TP53* mutant; missense R280K *TP53* mutation in DNA-binding domain, *BRCA1* wild-type) were treated with either TTFields at 150 khz, talazoparib at 2.5 µM or the combination for 24 hours. The combination showed an increase in PARP cleavage, p21(WAF1) and γH2AX upregulation compared to each treatment alone.

**Figure 3.**
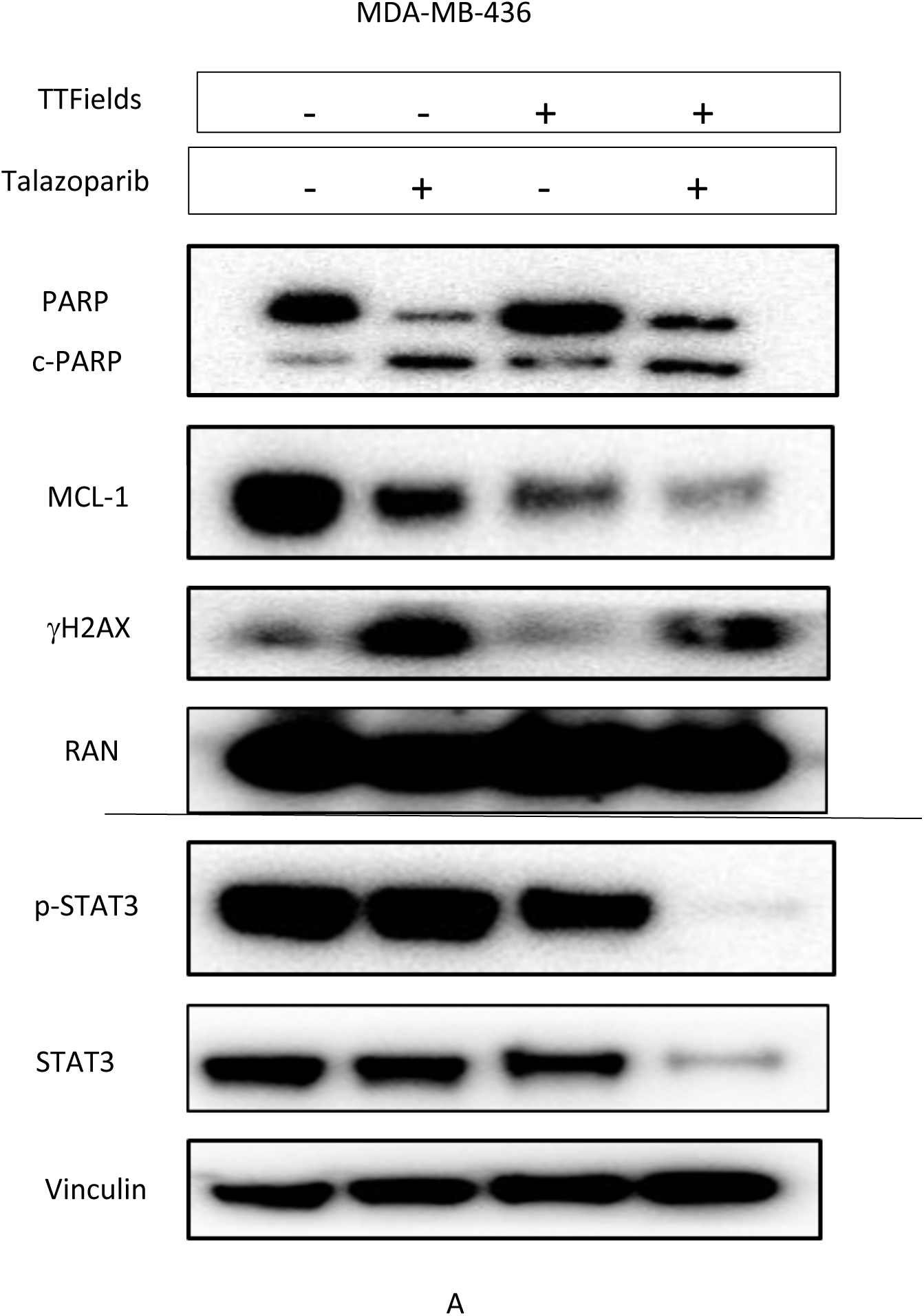
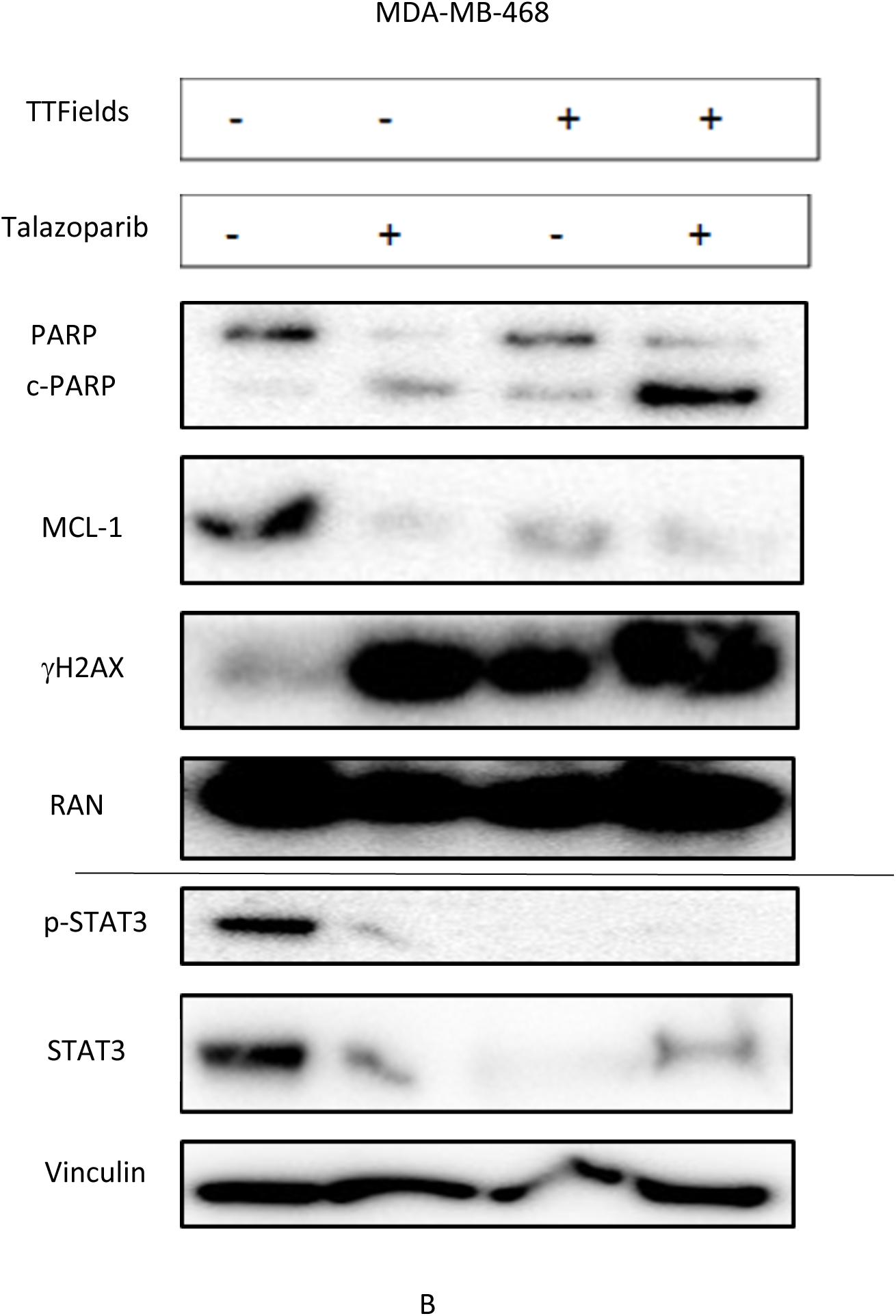
Enhanced cell death by TTFields -/+ talazoparib, independent of *TP53/BRCA1* mutational status, is associated with DNA damage, and reduced expression of pro-survival proteins including p-STAT3, STAT3, and MCL1. **(A)** Human MDA-MB-436 TNBC cells (*BRCA1* mutated) were treated with either TTFields at 150 khz, talazoparib at 2.5 µM or the combination for 72 hours. The combination and each Treatment caused PARP cleavage, p-STAT3, STAT3 and MCL-1 Downregulation (mild downregulation of p-STAT3, STAT3 and MCL-1 was observed following TTFields alone). Both talazoparib and TTFields induced DNA damage as evidenced by γH2AX upregulation. **(B)** The human MDA-MB-468 TNBC cells (*BRCA1* wild-type) were treated with either TTFields at 150 khz, talazoparib at 2.5 µM or the combination for 72 hours. There was more PARP cleavage, γH2AX with combination of TTFields plus talazoparib compared to each treatment alone. Both single treatments and combination of TTFields plus talazoparib reduced p-STAT3, STAT3 and MCL-1 expression in treated TNBC cells.

### Combination of PARP inhibitor talazoparib and TTFields led p-STAT3/STAT3, MCL-1 downregulation in TNBC cell lines

We further explored the underlying mechanisms by which TTFields and PARP inhibitor talazoparib reduced cell survival and induced cell death. MDA-MB-436 and MDA-MB-468 cells were treated with either TTFields, talazoparib or the combination for 72 hours. As mentioned earlier, STAT3 activates transcription of multiple cell growth factors. Activated p-STAT3 leads to MCL1 (pro-survival marker) transcription (25). We found that p-STAT3, STAT3, and MCL1 were decreased following exposure to TTFields together with talazoparib in the MDA-MB-436 and MDA-MB-468 TNBC cell lines (**Figure 3**). Further increase in the γH2AX marker of DNA damage and PARP cleavage was observed following exposure to the TTFields and talazoparib combination in the MDA-MB-468 cell line (**Figure 3B**).

### TTFields together with PARP inhibitor talazoparib reduces PDL-1 expression

Based on our findings showing an effect on STAT3, we further investigated PD-L1 expression, as previous studies have shown that STAT3 activation can induce PD-L1 upregulation (27). PDL-1 has multiple effects from immune escape to tumor progression (28). TTFields together with talazoparib Treatment in MDA-MB-231 and MDA-MB-436 led to PDL-1 downregulation after 24 hours of treatment (**Figure 4**).

**Figure 4.**
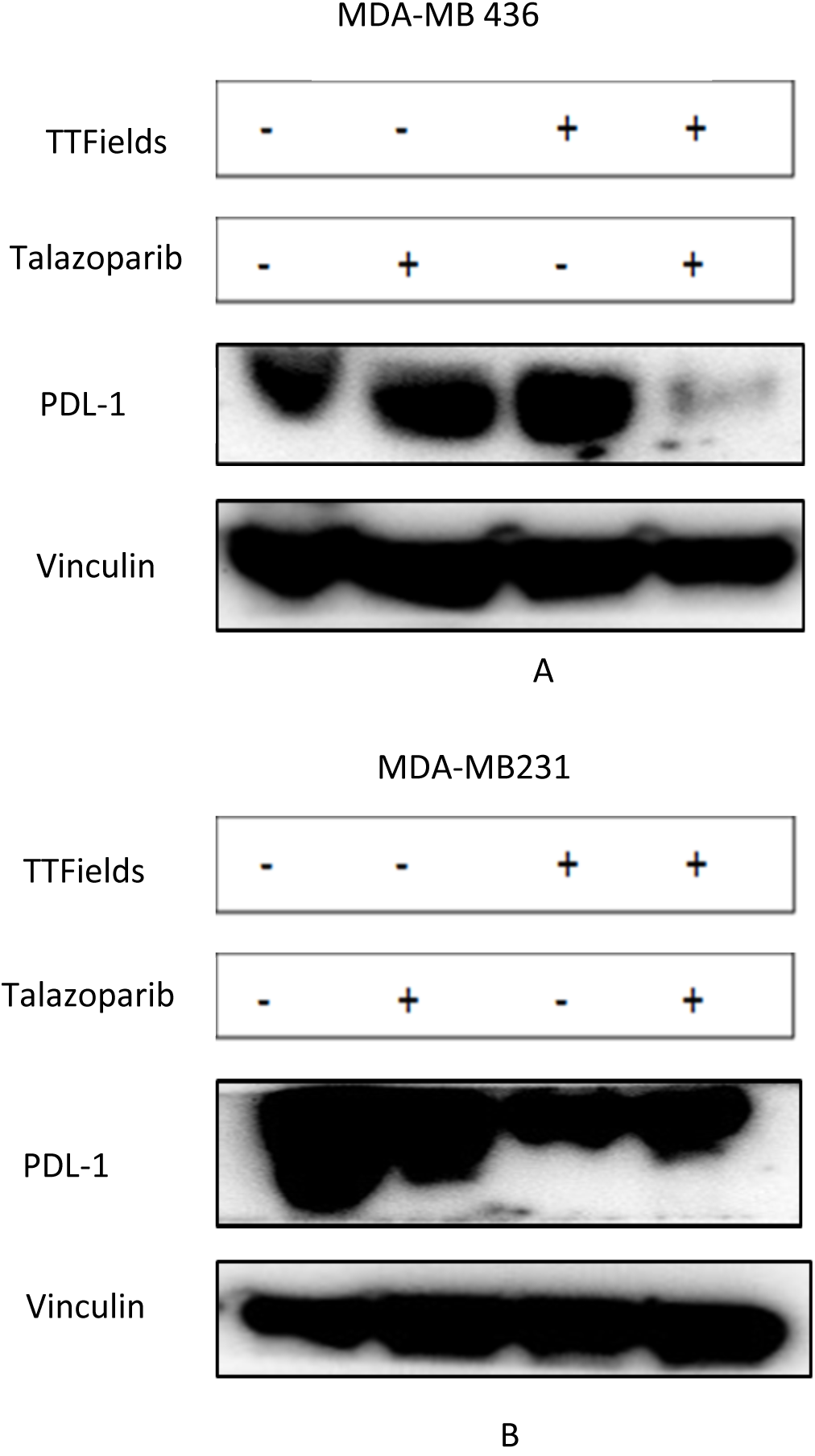
Reduced PDL-1 expression in TTFields and talazoparib-treated human TNBC cells occurs independently of *TP53/BRCA1* mutational status. **(A)** Human MDA-MB-436 TNBC cells (*BRCA1* mutated) were treated with either TTFields at 150 khz, talazoparib at 2.5 µM or the combination for 24 hours. PDL-1 protein expression was decreased in combination-treated TNBC cells. **(B)** Human MDA-MB-231 TNBC cells (*BRCA1* wild-type, *TP53* mutated) were treated with either TTFields at 150 khz, talazoparib at 2.5 µM or the combination for 24 hours. PDL-1 protein expression was decreased following TTFields and combination treatment. Milder decrease was observed following talazoparib treatment.

## Discussion

Treatment of triple-negative breast cancer (TNBC) presents significant challenges due to its high risk of relapse and resistance to chemotherapy. The development of effective new treatment strategies and novel therapeutic combinations is essential to improve patient outcomes (28). We investigated Tumor Treating Fields (TTFields) as a novel treatment option for breast cancer, exploring its cytotoxic effects and molecular mechanisms both alone and together with PARP inhibitor talazoparib.

Our results demonstrate that TTFields effectively downregulated key markers of cell growth, such as STAT3. Notably, talazoparib did not induce an increase in STAT3 activity as seen with Olaparib treatment, reported in previous studies. Additionally, the combination of TTFields and talazoparib further reduced STAT3 activity. The anti-apoptotic marker MCL-1 was significantly decreased both by TTFields alone and by the TTFields plus talazopraib combination.

The combination of TTFields plus talazopraib was effective in inducing cell death as evidenced by PARP cleavage and increased DNA damage, as indicated by elevated γH2AX levels. These effects were heterogeneous across different human TNBC cell lines. Furthermore, TTFields alone caused cell cycle dependent kinase inhibitor p21(WAF1/CDKN1A) protein upregulation, and the induction was increased by adding PARP inhibitor talazoparib. This observation aligns with previous studies describing the effect of TTFields on p21 in other cancer types (12,13,14,15,16). We demonstrate here that the induction of p21 is independent of *TP53* mutational status, suggesting a p53-independent mechanism.

TTFields together with PARP inhibitor talazoparib also caused PDL-1 downregulation. PD-L1 engagement with PD-1 on immune cells causes immune evasion in cancer immune surveillance (29). Since the interaction between PD-L1 on tumor cells and PD-1 on T cells can suppress T-cell activity, the decrease in PD-L1 observed in our study raises the possibility of reduced tumor-mediated immune suppression (30). PD-L1 is an important predictive biomarker for immune checkpoint inhibitor therapy in breast cancer, particularly in metastatic TNBC, but its predictive value can vary depending on the disease setting and the method used to assess PD-L1 expression (31). The implications of reduced PD-L1 by combined TTFields plus talazoparib for immune checkpoint inhibitor therapy remain unclear, as higher PD-L1 sensitizes T-cells to killing of tumor cells in the presence of immune checkpoint blockade therapy. However, the reduced PD-L1 by TTFields plus talazoparib may improve anti-tumor immunity by stimulating more endogenous (or adoptively transferred) T-cells to kill the tumor cells.

Previous studies have suggested that TTFields can affect the tumor immune response. TTFields have been shown to promote immunogenic cell death and immune cell activation, and their combination with anti-PD-1 treatment improved tumor control in preclinical models (32). In another *in vivo* study, TTFields together with immune checkpoint inhibitors, including anti-PD-L1 treatment, increased cytotoxic T-cell infiltration and improved tumor control compared with the individual treatments (33). These findings suggest that the relationship between TTFields, PD-L1 expression, and anti-tumor immunity may depend on the interaction between tumor cells and the surrounding immune microenvironment. Since our findings are based on tumor cells *in vitro*, further studies in immunocompetent mouse models are needed to determine whether the observed decrease in PD-L1 reduces tumor-mediated immune suppression and how it may affect response to PD-1/PD-L1-directed therapy.

In conclusion, our study demonstrates that TTFields, either alone or together with PARP inhibitor talazoparib, induces DNA damage, p53-independent upregulation of p21, reduces anti-apoptotic markers including p-STAT3, STAT3, and MCL1, reduces immune checkpoint PDL-1 protein expression and promotes cell death in human TNBC cells. Our preclinical results indicate that TTFields together with talazoparib may serve as effective therapeutic options in breast cancer.

## Acknowledgements

W.S.E-D. is an American Cancer Society Research Professor and is supported by the Mencoff Family University Professorship at Brown University. This work was supported by an AACR-Novocure grant to W.S.E-D. This work was previously presented in part at annual meeting of the American Association for Cancer Research (Ghandali, M., Srinivasan, P., Xia, Y., Wong, E.T., Sobol, R.W., Zhou, L., Carneiro, B.A., Graff, S.L., and El-Deiry, W.S. Tumor Treating Fields (TTFields) show efficacy in Triple-Negative breast cancer (TNBC) cells alone and in combination with PARP inhibitor Talazoparib. 115^th^ Annual AACR meeting, April 5-10, 2024, San Diego, CA.).

